# Probabilistic Graphical Models Relate Immune Status with Response to Neoadjuvant Chemo-Therapy in Breast Cancer

**DOI:** 10.1101/210112

**Authors:** Andrea Zapater-Moros, Angelo Gámez-Pozo, Guillermo Prado-Vázquez, Lucía Trilla-Fuertes, Jorge M Arevalillo, Mariana Díaz-Almirón, Hilario Navarro, Paloma Maín, Jaime Feliú, Pilar Zamora, Enrique Espinosa, Juan Ángel Fresno Vara

**Affiliations:** Molecular Oncology & Pathology Lab, Institute of Medical and Molecular Genetics-INGEMM, La Paz University Hospital-IdiPAZ, Madrid, Spain.; Biomedica Molecular Medicine SL, Madrid, Spain.; Operational Research and Numerical Analysis, National Distance Education University (UNED), Ma-drid, Spain.; Biostatistics Unit. La Paz University Hospital-IdiPAZ, Madrid, Spain.; Department of Statistics and Operations Research, Faculty of Mathematics, Complutense University of Madrid, Madrid, Spain.; Medical Oncology Service, La Paz University Hospital-IdiPAZ, Madrid, Spain.; CIBERONC

**Author notes:** Corresponding Author Dr. Juan Angel Fresno Vara Hospital Universitario La Paz-IdiPAZ Paseo de la Castellana, 261 28046 Madrid. Competing interests JAFV, EE and AG-P are shareholders in Biomedica Molecular Medicine SL. LT-F is an employee of Biomedica Molecular Medicine SL. The other authors declare no potential conflicts of interest.

**Keywords:** breast cancer, crowding, neoadjuvant chemotherapy, molecular subtypes, probabilistic graphical models, immune status

## Abstract

Breast cancer is the most frequent tumor in women and its incidence is increasing. Neoadjuvant chemotherapy has become standard of care as a complement to surgery in locally advanced or poor-prognosis early stage disease. The achievement of a complete response to neoadjuvant chemotherapy correlates with prognosis but it is not possible to predict who will obtain an excellent response. The molecular analysis of the tumor offers a unique opportunity to unveil predictive factors. In this work, gene expression profiling in 279 tumor samples from patients receiving neoadjuvant chemotherapy was performed and probabilistic graphical models were used. This approach enables addressing biological and clinical questions from a Systems Biology perspective, allowing to deal with large gene expression data and their interactions. Tumors presenting complete response to neoadjuvant chemotherapy had a higher activity of immune related functions compared to resistant tumors. Similarly, samples from complete responders presented higher expression of lymphocyte cell lineage markers, immune-activating and immune-suppressive markers, which may correlate with tumor infiltration by lymphocytes (TILs). These results suggest that the patient’s immune system plays a key role in tumor response to neoadjuvant treatment. However, future studies with larger cohorts are necessary to validate these hypotheses.

## INTRODUCTION

Breast cancer is the most common neoplasm and the fifth cause of cancer-associated death among women (1). Estrogen receptor (ER), progesterone receptor (PR), and human epidermal growth factor receptor 2 (HER2) provide a system of classification and clinical diagnosis. Seventy percent of the tumors are hormonal receptor positive, and HER2 overexpression is observed in 15% of cases. ER+ and PR+ tumors respond to endocrine therapy, whereas tumors overexpressing HER2 respond to targeted therapies such as trastuzumab (2, 3). Tumors negative for ER, PR and HER2 are known as Triple Negative Breast Cancer (TNBC) and do not respond to the aforementioned therapies.

A molecular classification of breast cancer defined four intrinsic subtypes (4). Luminal A disease, which accounts for 67% of all tumors, shows high expression of genes related to hormone receptors and low expression of genes related to cell proliferation. Luminal B, HER2-enriched and Basal-like subtypes have a more aggressive phenotype (5) (6, 7).

Neoadjuvant chemotherapy has been increasingly administered to reduce the size of primary tumor, thus increasing the likelihood of breast conservation and enhancing survival (8). Currently, there is no clinically useful molecular predictor of response to cytotoxic drugs in the neoadjuvant setting. Clinical parameters or the expression of single molecular markers (ie, Bcl-2, p53, MDR-1, and so on) show weak association with response and are not regimen-specific. Molecular subtyping may offer some help, as Luminal B and Basal-like tumors respond better than Luminal A tumors (9), but this is not accurate enough to make clinical decisions. As a consequence, many patients suffer the toxicity of useless neoadjuvant chemotherapy.

The objective of this study was to evaluate differences in gene expression patterns of breast cancer tumors from patients who had undergone neoadjuvant chemotherapy through a Systems Biology perspective. This study has been carried out using probabilistic graphical models, providing insights into the molecular biology of tumor response, allowing its use as a predictive model for response. These statistically inferred networks provide a deeper level of biological understanding in two main directions: giving support to previously identified biological observations, and giving new insights regarding novel biological interactions. Moreover, the transcriptional network approach has proven to be useful to unveil transcriptional regulation in breast cancer (10, 11).

## MATERIALS & METHODS

### Patient’s and samples origin and characteristics

A breast cancer tumor dataset, including gene expression and clinical data, was obtained from the Gene Expression Omnibus (GSE41998) and from a phase II trial (NCT00455533). Gene expression profiling was measured using an Affymetrix GeneChip, normalized and log2 transformed. Surgical specimens were evaluated by a pathologist at each study site. The pathological response was evaluated as the primary endpoint. A pathological complete response (CR) was defined by no histologic evidence of residual invasive adenocarcinoma in the breast and axillary lymph nodes, with or without the presence of ductal carcinoma in situ (12). Responses were categorized as complete, partial, stabilization or progression.

### Gene expression data preprocessing

The PAM50 method was used as previously described to assign a molecular subtype to each sample (13). Lehmann subtypes for TNBC were assigned in two steps (14). First, samples were assigned to Lehmann’s seven subtypes using centroids constructed from 77 previously assigned tumors in GSE31519 dataset. Then, the IM and MSL groups were redefined as previously described (14).

### Probabilistic graphical models construction

A functional structure was developed using undirected probabilistic graphical models (PGMs) as previously described (10). Briefly, PGMs compatible with high-dimensionality were chosen. The result is an undirected graphical model with local minimum Bayesian Information Criterion. DAVID 6.8 was used to assign a biological function to each node in the networks, using “homo sapiens” as background list and selecting only GOTERM-FAT and Biocarta and KEGG pathways categories. Functional activity of each node was calculated by the mean expression of the genes in each node. To visualize node activities, data from all tumors used to construct the network were mean centred prior to its inclusion into the network.

### Statistics and software suites

Differences between groups were assessed using Kruskal-Wallis test, Mann-Whitney test and Dunn`s multiple comparisons test using GraphPad Prism 6. Box-and-whisker plots are Tukey boxplots. All p-values were two-sided and p<0.05 was considered statistically significant. Ordinal logistic regression analysis was performed in SAS using logistic procedure. Network analyses were performed in MeV and Cytoscape 3.2.1 software suites.

## RESULTS

### Patient’s characteristics

279 patients with histologically-confirmed primary non-metastatic breast adenocarcinoma were included. They all had untreated tumors of at least 2 cm in size (T2–3, N0– 3) regardless of hormone receptor or HER2 expression status. Clinical data were obtained from phase II trial (NCT00455533). Patient’s clinical characteristics are provided in Table 1. On the basis of ER, PR and HER2 status, 111 tumors patients (39.78%) were ER+ or PR+ and Her2-(ER+ for now on), 28 (10.04%) were HER2+ and 140 (50.18%) were classified as TNBC. Patients received sequential neoadjuvant therapy starting with 4 cycles of doxorubicin/cyclophosphamide (AC), followed by 1:1 randomization to either ixabepilone or paclitaxel. All patients underwent definitive breast surgery 4 to 6 weeks after the last dose of ixabepilone or paclitaxel, consisting of either a lumpectomy with axillary dissection or modified radical mastectomy. Regarding pathological response, 40 (14.34%) patients achieved a complete response (CR), 161 (57.71%) achieved a partial response (PR), 64 (22.94%) had stable disease (SD) and 5 (1.79%) had progressive disease (PD).

### Molecular stratification of tumors

Molecular subtypes were defined by PAM50 assignment (13). Of the initial 279 patients, 116 (41.58%) patients were classified as Basal-like subtype, 15 patients (5.38%) as HER2+, 66 (23.66%) as Luminal A, 62 (22.22%) as Luminal B, and 15 (5.38%) as Normal-like. Five patients could not be assigned. A sub-classification of TNBCs was performed based on Lehmann’s classification as previously described (14); 25 (8.96%) TNBC tumors were Basal-like 1, 83 (29.75%) Basal-like 2 subgroup, 6 (2.15%) Luminal Androgen Receptor, and 26 (9.32%) Mesenchymal.

### Breast cancer systems biology

Gene expression data from all tumor samples were used to build a probabilistic graphical model, with no other *a priori* information. The resulting graph was divided in eighteen branches (functional nodes) and a main function was assigned to each node by gene ontology analysis. The structure of the probabilistic graphical model clearly reflected different biological functions. Functional node activities were then calculated as previously showed (10, 15) (Figure 1).

**Fig. 1.**
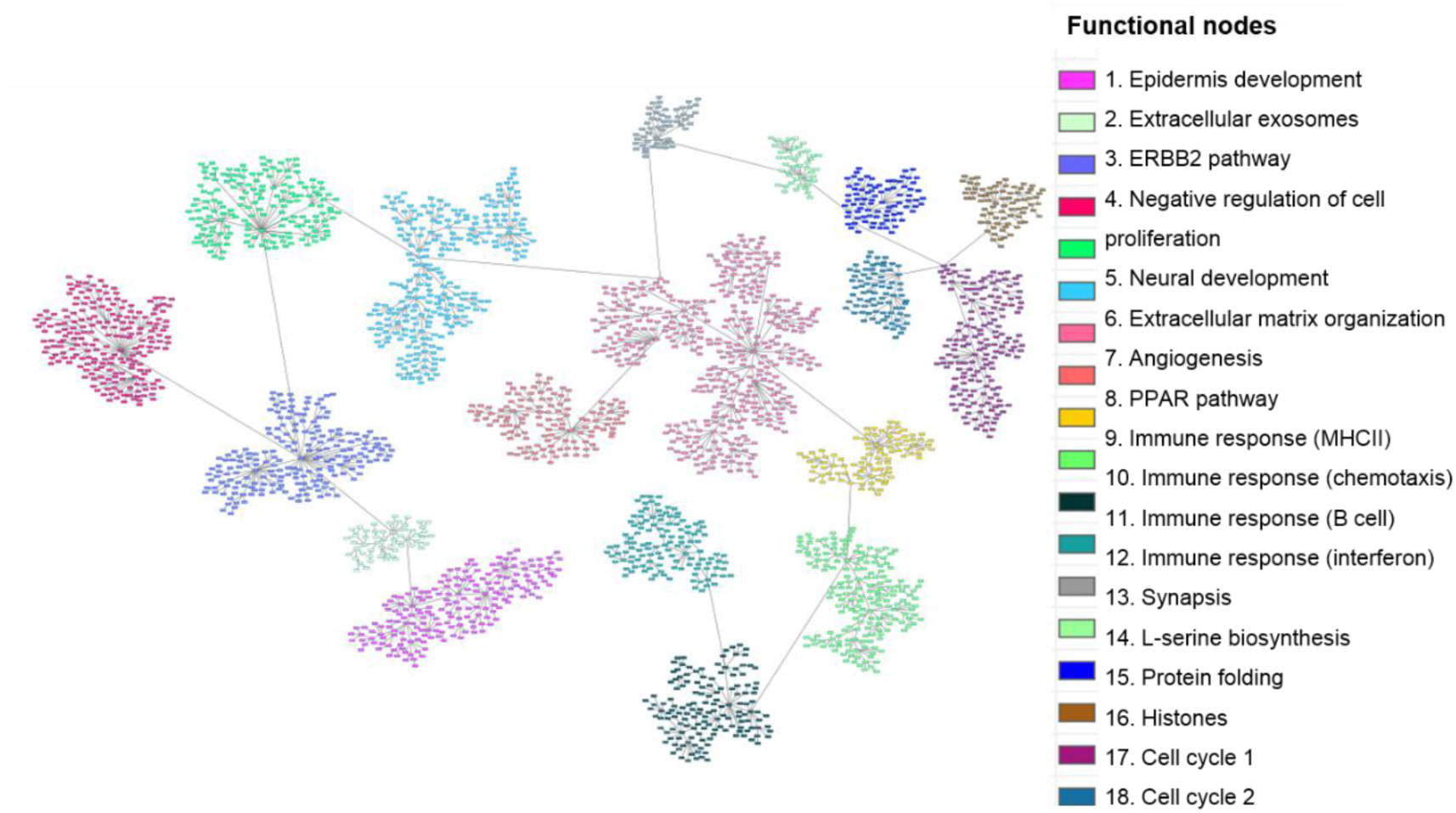
Breast cancer network. Probabilistic graphical model from 279 tumors gene expression data divided in eighteen functional nodes harboring one or two predominant biological functions. Each node (box) represents one gene and each grey line (edges) connects genes with correlated expression.

### Functional structure of response to neoadjuvant chemotherapy

Patients were classified according to pathological response regardless of their tumor molecular subtype to study the response to neoadjuvant chemotherapy. Significant differences between functional node activities were observed in “Immune response (MHCII)” (9), “Immune response (B cell)” (11) and “Immune response (Interferon)” (12) nodes, in which, tumors attaining a complete response had higher activation (Figure 2). A progressive decrease in the activity of immune functional nodes was seen depending on the response, being higher in tumors obtaining a CR and absent in those having a progression. Additionally, the relationship of immune nodes activities with the pathological response was evaluated using an ordinal logistic regression analysis. This analysis revealed that an increment of one unit in node 9, 11 and 12 activities increased the probability of a favorable response 1.739, 1.435 and 1.629 times respectively. By contrast, one unit increase in the activity of node 10 increased 0.519 times the probability of having an unfavorable response. On the other hand, PD tumors showed higher functional activity in “Cell cycle 1" (17) and “Cell cycle 2" (18) nodes, followed by CR tumors.

**Figure 2.**
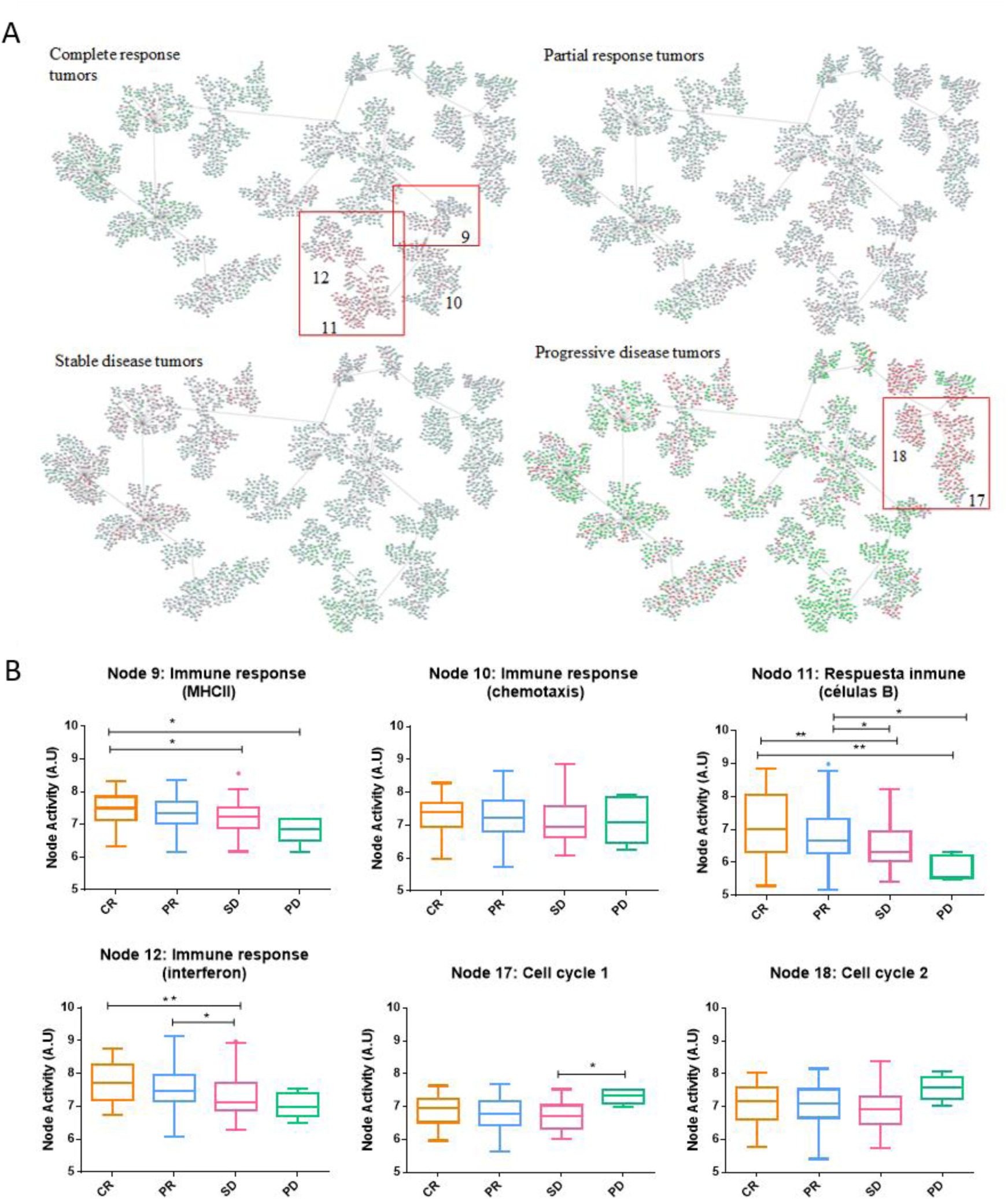
Breast cancer network by pathological response groups. Detail of nodes with the highest activity in each of the subgroups. Genes with an expression below 0 were represented in green; genes with an expression around 0 were represented in grey and genes with an expression above zero were represented in red. Functional node activities differences between pathological response groups: Box-and-whisker plots are Tukey boxplots. All p-values were two-sided and p<0.05 was considered statistically significant. P-value < 0.05 (*); p-value < 0.01(**). A.U: arbitrary units.

### Functional characterization of molecular subtypes

Patients in the network were further classified according to their molecular subtype (Basal-like, Luminal A, Luminal B, HER2+ and Normal-like). Basal-like tumors were also classified according to Lehmann's subtypes. “Immune response (MHCII)” (9) node activity was higher in Luminal A and Normal-like subtypes while Basal-like tumors showed higher functional node activity in “Immune response (chemotaxis)” (10), “Immune response (B cell)” (11) and “Immune response (Interferon)” (12), as well as in “Cell cycle 1 and 2” (17 and 18) nodes (Figure 3).

**Figure 3.**
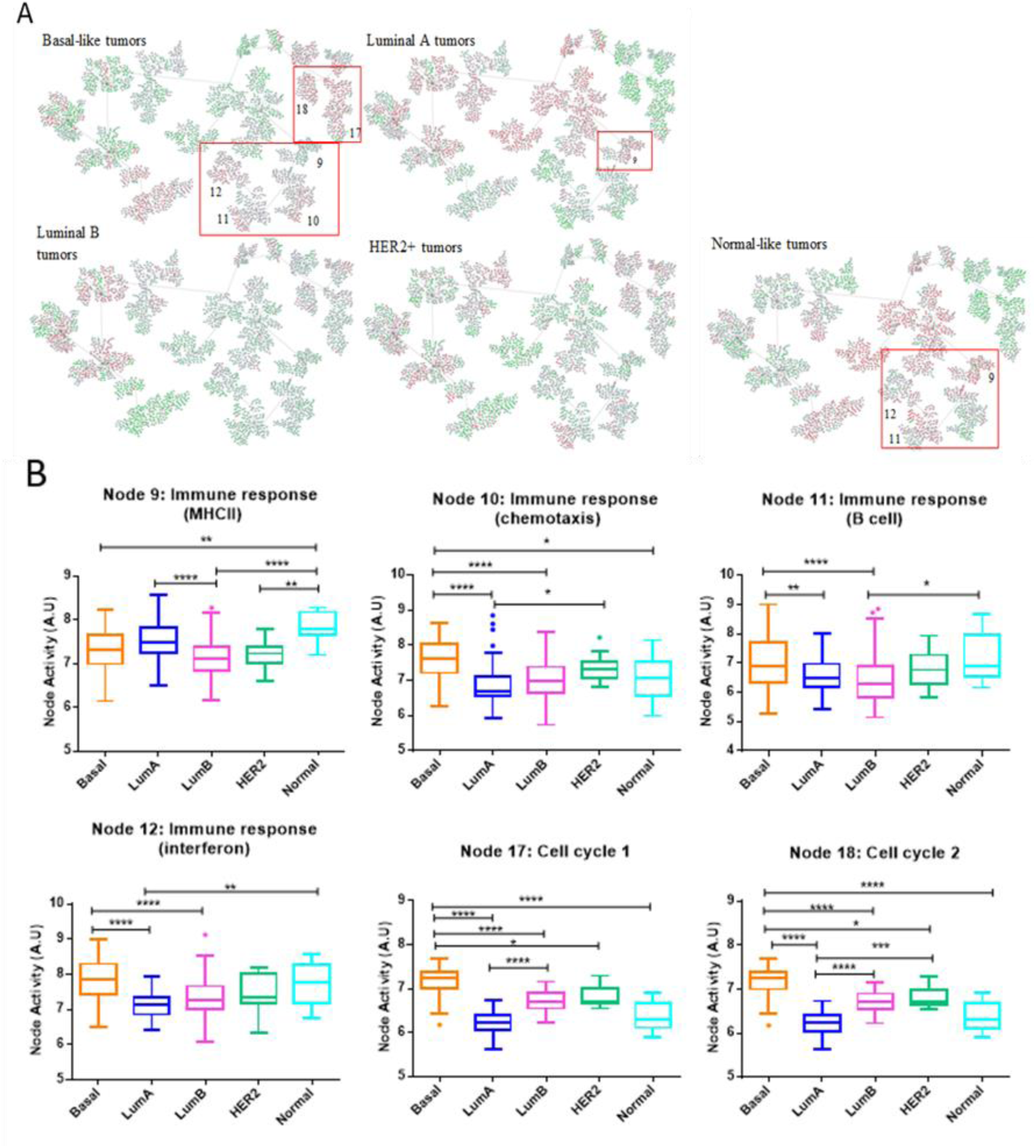
Breast cancer network by breast cancer molecular subtypes. A) Detail of nodes with the highest activity in each of the subgroups. Genes with an expression below 0 were represented in green; genes with an expression around 0 were represented in grey and genes with an expression above zero were represented in red. B) Functional node activities differences between molecular subtypes: Box-and-whisker plots are Tukey boxplots. All p-values were two-sided and p<0.05 was considered statistically significant. P-value < 0.05 (*); p-value < 0.01(**); P-value< 0.001 (***); P-value <0.0001 (****). A.U: arbitrary units.

Concerning TNBCs sub-classification, BL2 subtype showed a higher functional activity in “Immune response” (9, 10, 11 and 12) nodes whereas it was observed higher “Cell cycle 1 and 2” (17 and 18) nodes activities in BL1 tumors

In order to evaluate the functional implications between molecular subtypes and response to neoadjuvant therapy, data from patients of the same molecular subtype were mean centred and analysed independently. Luminal A group included no PD, whereas only one PD was found in Luminal B group, and was excluded from this analysis. Normal-like and HER2+ tumors were insufficient to perform subsequent analyses.

Concerning Luminal A and Luminal B subtypes, “Immune response (MHCII)” (9), "Immune response (chemotaxis)" (10), “Immune response (B cell)” (11) and “Immune response (Interferon)” (12) functional nodes activities were higher in tumors attaining a CR, although differences were not statistically significant. As in the case of Basal-like tumors, a progressive decrease of activity in these nodes was observed from CR to SD.

In Basal-like subtype, tumors attaining a CR showed significant differences in “Immune response (B cell)” (11) and “Immune response (Interferon)” (12) nodes activities while “Immune response (MHCII)” (9) node activity was higher in tumors showing a PR. PD tumors showed a higher functional node activity in Cell cycle 1" (17) and Cell cycle 2" (18). Regarding TNBC, in the BL1 subtype, the activity of nodes “Immune response (B cell)” (11) and “Immune response (Interferon)” (12) was higher in CR than in PR. “Immune response (MHCII)” (9), "Immune response (chemotaxis)" (10) node activities also were higher in CR tumors but without statistics differences. In the BL2 subtype, CR tumors showed a significant higher activity in “Immune response (MHCII)” (9) and “Immune response (B cell)” (11) nodes. However, SD tumors showed a higher activity in "Immune response (chemotaxis)" (10) node. In Mesenchymal subtype, CR tumors showed higher activity in “Immune response (MHCII)” (9) and “Immune response (B cell)” (11) nodes but without being statistically significant.

Three separate probabilistic graphical models were built for Basal-like, Luminal-A and Luminal-B subtypes. As in the global network, tumors experiencing a CR had an increased activity of immune response-related nodes, although differences were not statistically significant.

### Immunological markers

Markers of tumor-infiltrating lymphocytes were analysed to further characterize response according to the immune status. For that, patients were separated according to their pathological response and marker gene expression levels were compared between groups. Tumors obtaining a CR showed significantly higher expression levels of cell lineage markers (CD4, CD8 and CD20) as well as immune-activating (IGKC, CXCL9, CCL5, CXCL13) and immunosuppressive markers (IDO1, PD1) compared to the rest of tumors (Figure 4).

**Figure 4.**
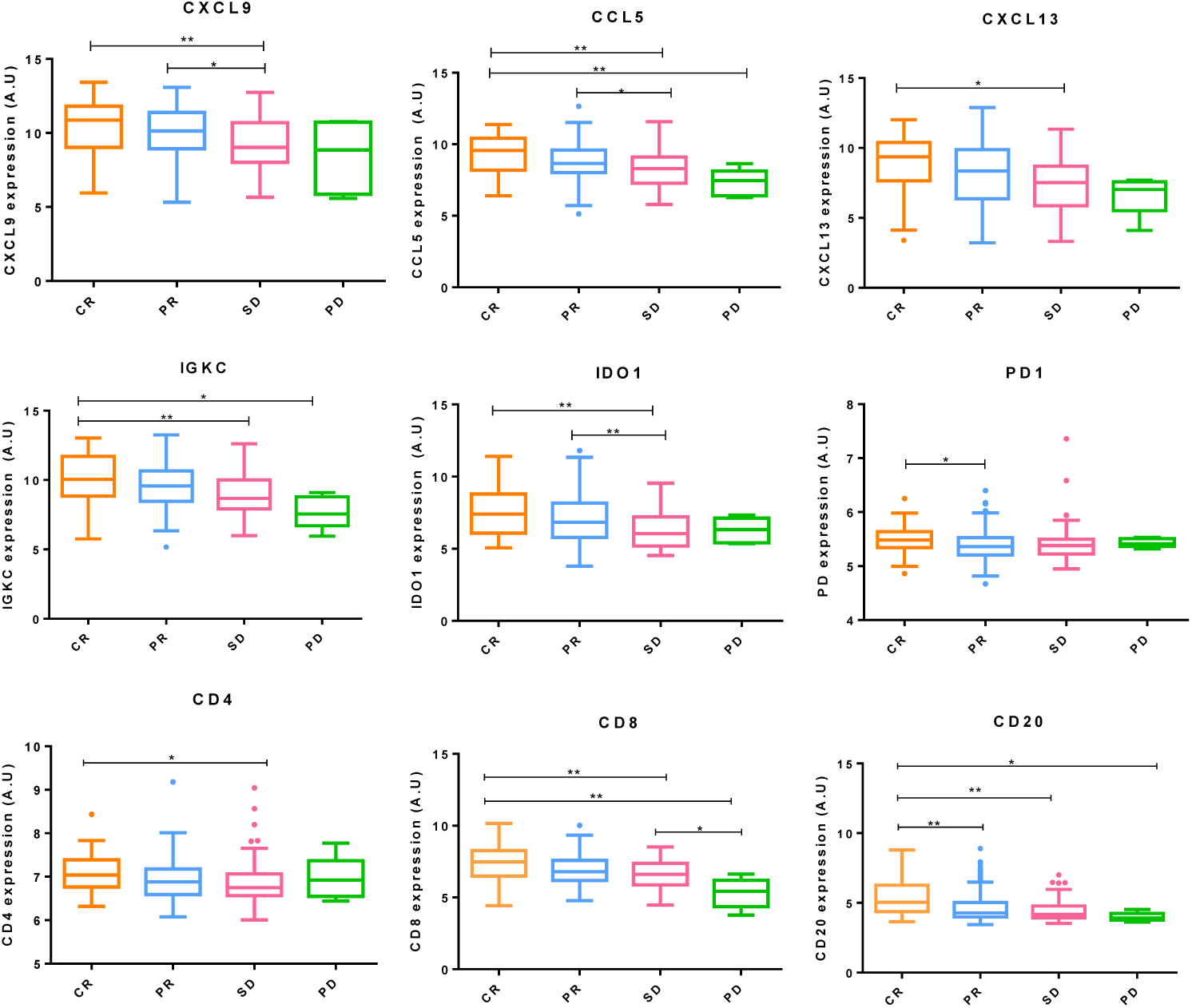
Immunological markers expression. Immune-activating, immunosuppressive and cell lineage markers gene expression between pathological response groups. Box-and-whisker plots are Tukey boxplots. All p-values were two-sided and p<0.05 was considered statistically significant. P-value < 0.05 (*); p-value < 0.01(**); p-value < 0.001 (***); p-value <0.0001 (****). A.U: arbitrary units.

## DISCUSSION

In this work, a gene expression-based probabilistic graphical model analysis of breast cancer showed that immune functional nodes were related to pathological response to neoadjuvant chemotherapy. This correlation did not depend on the molecular subtype, indicating that a Systems Biology approach complements knowledge obtained from other research methods. Non-directed probabilistic graphical models allow managing large gene datasets and underscoring lots of gene interactions, many of which have not been previously described.

The activity of immune nodes was higher in tumors attaining a CR and decreased with the intensity of response. Tumors progressing on chemotherapy also showed increased activity in the nodes "Cell division 1" (17) and "Cell division 2" (18). These results suggest that the patient's immune system plays a crucial role in the response to neoadjuvant chemotherapy. Previous studies suggest that conventional therapies are effective in patients exhibiting some degree of immune activation in the tumor (16), supporting our findings. Chemotherapy may mediate the “immunogenic” death of tumor cells, thus facilitating an immune response against the disease (17).

As expected, all tumors attaining a CR-regardless of molecular subtype-showed significantly higher levels of cell lineage markers (CD4, CD8 and CD20) as well as immune-activating (IGKC, CXCL9, CCL5, CXCL13) and immunosuppressive markers (IDO1, PD1), suggesting a greater infiltration of immune cells High tumor-infiltrating lymphocytes (TILs) levels have been associated with increased CR rates in ER+ HER2+/-tumors (18) and also in TNBC (19). However, high levels of PD-1+ TILs or Foxp3+ TILs have been related with poor prognosis (18). Therefore, immune cell subpopulation profiles could help to predict response to neoadjuvant chemotherapy.

Basal-like and HER2+ subgroups have been associated with highest CR rates as opposed to Luminal and Normal-like tumors (20, 21). In the present cohort, Basal-like tumors had the highest CR rate as expected. However, the CR rate was poor in HER2+ tumors, probable because these patients did not receive HER2 targeted therapy. Also as expected, BL1 tumors achieved a CR more commonly than TNBC subtypes (22). Although node "Immune response (MHCII)" (9) had higher activity in Luminal A and Normal subtypes, the remaining nodes related to immune response were more active in Basal-like tumors. Basal-like also had the highest activity in the node "Cell cycle" (17 and 18), which is in accordance with the fact proliferation renders tumor cells more sensitive to chemotherapy (6).

The neoadjuvant setting is appealing in the field of drug development because it allows early evaluation of efficacy. However, not all patients benefit from this approach, so markers predicting response to neoadjuvant chemotherapy are clearly needed. Our results suggest that immune activation in the tumor may identify responders. Although validation is needed, the use of these markers can help to determine the future use of neoadjuvant chemotherapy in breast cancer.

## Notes

Funding statement This study was supported by Instituto de Salud Carlos III, Spanish Economy and Competitiveness Ministry, Spain and co-funded by the FEDER program, “Una forma de hacer Europa” (PI15/01310). LT-F is supported by the Spanish Economy and Competitiveness Ministry (DI-15-07614). The funders had no role in the study design, data collection and analysis, decision to publish or preparation of the manuscript.

## BIBLIOGRAPHY

1. Ferlay J, Soerjomataram I, Dikshit R, Eser S, Mathers C, Rebelo M, et al. Cancer incidence and mortality worldwide: sources, methods and major patterns in GLOBOCAN 2012. Int J Cancer. 2015;136(5):E359–86. Epub 2014/10/09. doi: 10.1002/ijc.29210. PubMed PMID: 25220842.

2. Dawson SJ, Rueda OM, Aparicio S, Caldas C. A new genome-driven integarated classification of breast cancer and its implications. The EMBO journal. 2013;32(5):617–28.

3. Honrado E, Benítez J, Palacios J. The pathology of hereditary breast cancer. Hered Cancer Clin Pract. 2004;2(3):131–8. Epub 2004/07/15. doi: 10.1186/1897-4287-2-3-131. PubMed PMID: 20233467; PubMed Central PMCID: PMCPMC4392521.

4. Perue C, Sorlie T, Elsen M, van de Rijn M, Jeffrey S, Rees C, et al. Molecular portraits of human breast tumors. Nature. 2000;406(6797):747–52.

5. Kennecke H, Yerushalmi R, Woods R, Cheang MCU, Voduc D, Speers CH, et al. Metastatic behavior of breast cancer subtypes. Journal of clinical oncology. 2010;28(20):3271–7.

6. van de Vijver MJ, He YD, van't Veer LJ, Dai H, Hart AA, Voskuil DW, et al. A gene-expression signature as a predictor of survival in breast cancer. N Engl J Med. 2002;347(25):1999–2009. doi: 10.1056/NEJMoa021967. PubMed PMID: 12490681.

7. Badve S, Dabbs DJ, Schnitt SJ, Baehner FL, Decker T, Eusebi V, et al. Basal-like and triple-negative breast cancers: a critical review with an emphasis on the implications for pathologists and oncologists. Modern Pathology. 2011;24(2):157–67.

8. Smith IC, Heys SD, Hutcheon AW, Miller ID, Payne S, Gilbert FJ, et al. Neoadjuvant chemotherapy in breast cancer: significantly enhanced response with docetaxel. Journal of Clinical Oncology. 2002;20(6):1456–66.

9. Eroles P, Bosch A, Pérez-Fidalgo JA, Lluch A. Molecular biology in breast cancer: intrinsic subtypes and signaling pathways. Cancer treatment reviews. 2012;38(6):698–707.

10. Gámez-Pozo A, Berges-Soria J, Arevalillo JM, Nanni P, López-Vacas R, Navarro H, et al. Combined label-free quantitative proteomics and microRNA expression analysis of breast cancer unravel molecular differences with clinical implications. Cancer research. 2015;75(11):2243–53.

11. de Anda-Jáuregui G, Velázquez-Caldelas TE, Espinal-Enríquez J, Hernández-Lemus E. Transcriptional Network Architecture of Breast Cancer Molecular Subtypes. Frontiers in Physiology. 2016;7.

12. Horak CE, Pusztai L, Xing G, Trifan OC, Saura C, Tseng L-M, et al. Biomarker analysis of neoadjuvant doxorubicin/cyclophosphamide followed by ixabepilone or Paclitaxel in early-stage breast cancer. Clinical Cancer Research. 2013;19(6):1587–95.

13. Parker JS, Mullins M, Cheang MC, Leung S, Voduc D, Vickery T, et al. Supervised risk predictor of breast cancer based on intrinsic subtypes. Journal of clinical oncology. 2009;27(8):1160–7.

14. Lehmann BD, Jovanović B, Chen X, Estrada MV, Johnson KN, Shyr Y, et al. Refinement of triple-negative breast cancer molecular subtypes: implications for neoadjuvant chemotherapy selection. PLoS One. 2016;11(6):e0157368.

15. Gámez-Pozo A, Trilla-Fuertes L, Berges-Soria J, Selevsek N, López-Vacas R, Díaz-Almirón M, et al. Functional proteomics outlines the complexity of breast cancer molecular subtypes. Scientific Reports. 2017;7(1):10100.

16. Denkert C, Loibl S, Noske A, Roller M, Müller BM, Komor M, et al. Tumor-associated lymphocytes as an independent predictor of response to neoadjuvant chemotherapy in breast cancer. Journal of clinical oncology. 2009;28(1):105–13.

17. Ghiringhelli F, Apetoh L. The interplay between the immune system and chemotherapy: emerging methods for optimizing therapy. Expert review of clinical immunology. 2014;10(1):19–30.

18. Yu X, Zhang Z, Wang Z, Wu P, Qiu F, Huang J. Prognostic and predictive value of tumor-infiltrating lymphocytes in breast cancer: a systematic review and meta-analysis. Clinical and Translational Oncology. 2016;18(5):497–506.

19. Denkert C, Von Minckwitz G, Brase JC, Sinn BV, Gade S, Kronenwett R, et al. Tumor-infiltrating lymphocytes and response to neoadjuvant chemotherapy with or without carboplatin in human epidermal growth factor receptor 2–positive and triple-negative primary breast cancers. Journal of clinical oncology. 2014;33(9):983–91.

20. Kuerer HM, Newman LA, Smith TL, Ames FC, Hunt KK, Dhingra K, et al. Clinical course of breast cancer patients with complete pathologic primary tumor and axillary lymph node response to doxorubicin-based neoadjuvant chemotherapy. Journal of Clinical Oncology. 1999;17(2):460–.

21. Rouzier R, Perou CM, Symmans WF, Ibrahim N, Cristofanilli M, Anderson K, et al. Breast cancer molecular subtypes respond differently to preoperative chemotherapy. Clinical Cancer Research. 2005;11(16):5678–85.

22. Masuda H, Baggerly KA, Wang Y, Zhang Y, Gonzalez-Angulo AM, Meric-Bernstam F, et al. Differential response to neoadjuvant chemotherapy among 7 triple-negative breast cancer molecular subtypes. Clinical Cancer Research. 2013;19(19):5533–40.

